# Interactive effects of temperature and salinity on metabolism and activity of the copepod *Tigriopus californicus*

**DOI:** 10.1101/2024.05.17.594749

**Authors:** Caroline E. Terry, Josie A. Liebzeit, Ella M. Purvis, W. Wesley Dowd

## Abstract

In natural environments two or more abiotic parameters often vary simultaneously, and interactions between covarying parameters frequently result in unpredictable, non-additive biological responses. To better understand the mechanisms and consequences of interactions between multiple stressors it is important to study their effects on both survival and performance. The splashpool copepod *Tigriopus californicus* tolerates extremely variable abiotic conditions and exhibits a non-additive, antagonistic interaction resulting in higher survival when simultaneously exposed to high salinity and acute heat stress. Here, we investigated *T. californicus’* response in activity and oxygen consumption under simultaneous manipulation of salinity and temperature to identify if this interaction also arises in these sublethal measures of performance. Oxygen consumption and activity rates decreased with increasing assay salinity. Oxygen consumption also sharply increased in response to acute transfer to lower salinities, an effect that was absent upon transfer to higher salinities. Elevated temperature led to reduced rates of activity overall, resulting in no discernible impact of increased temperature on routine metabolic rates. This suggests that swimming activity has a non-negligible effect on copepod’s metabolic rates and must be accounted for in metabolic studies. Temperature also interacted with assay salinity to affect activity and with acclimation salinity to affect routine metabolic rates upon acute salinity transfer, implying that the sublethal impacts of these co-varying factors are also not predictable from experiments that study them in isolation.

**Summary Statement:** Temperature and salinity interact to affect metabolic rate in the copepod *Tigriopus californicus*, but the stressors’ individual effects and their interaction are complicated by concurrent changes in activity.

## Introduction

Natural environments host a multitude of abiotic factors that can pose physiological challenges for organisms. These factors often covary, requiring organisms to respond to changes in multiple stressors at once (Denny and Dowd, 2022; Morris and Taylor, 1983). Marine environments are no exception, presenting various abiotic pressures from changes in temperature, oxygen concentration, pH, salinity, and more. Furthermore, global climate change is exacerbating the effects of these stressors, leading to rising temperatures, ocean acidification, increasingly hypoxic conditions, and increases in magnitude, frequency, duration, and variability of extreme abiotic events (Crain et al., 2008; Jentsch et al., 2007). Intensification of each of these stressors due to climate change is not happening in isolation, and variation in multiple stressors simultaneously can sometimes lead to compounding effects and additional costs on organisms (Crain et al., 2008; Sokolova, 2013). However, in lab experiments, these stressors are most often studied in isolation.

An organism’s response to an individual stressor is not necessarily reflective of responses in a natural environment, where multiple abiotic factors demand concurrent behavioral or physiological adjustments. Indeed, when faced with multiple stressors, organismal responses in survival and performance can be antagonistic or synergistic, rather than the strictly additive response that might be predicted from single-stressor exposures (Cline et al., 2020; Crain et al., 2008; deMayo et al., 2021; Kelly et al., 2016; Orr et al., 2022; Todgham and Stillman, 2013). In an additive scenario, the response to a combination of stressors would be equal to the sum of their individual effects. In non-additive scenarios, the effect of multiple stressors is lower (antagonistic) or greater (synergistic) in magnitude than the sum of those stressors’ individual effects. Non-additive interactions may comprise the majority of interaction outcomes, yet our ability to predict these patterns falls short (Crain et al., 2008). For example, the intertidal crab *Pachygrapsus marmoratus* demonstrates an increase in thermal tolerance when exposed to acute heat stress under more acidic conditions compared to heat-stressed crabs under normal pH (Madeira et al., 2014). Because of the prevalence of such non-additive interactions, investigating physiological effects of changing environments using mostly single-factor studies can miss important multi-stressor interactions, limiting our potential to make inferences about natural populations.

Measures of mortality and fitness at the individual and population level are often used as indicators of the impact of environmental stress, especially after extreme climate events. These responses are also most frequently measured when investigating responses to multiple environmental stressors (Orr et al., 2020). However, many effects of environmental stress are not so apparent. Instead, they are driven by sublethal responses, whose repercussions may be more subtle and take more time to play out (Pan et al., 2015). As such, measures of survival and fitness do not capture the whole picture of stress responses, necessitating complementary investigation of other response metrics. The manifestation of multi-stressor interactions at sublethal levels (e.g., performance, behavior) can have profound long-term implications, ultimately affecting survival, ecological interactions, and adaptive potential in the face of climate change (Orr et al., 2022; Pan et al., 2015). For example, metabolic rate responses to environmental change can indicate the severity of stress experienced (McAllen et al., 1999). They can also provide insights into how energy is allocated to homeostatic biochemical responses (e.g., protein turnover, ion transport), energy that is consequently unavailable for foraging, growing, reproducing, or responding to other stressors (Pan et al., 2015; Pörtner, 2010; Sokolova, 2013). Similarly, an organism’s activity levels can have important impacts on its success in predator avoidance, mate-finding, foraging, or behaviorally adjusting to environmental stress (Dinh et al., 2020; Riddell et al., 2005; Sloman et al., 2008; Speers-Roesch et al., 2018). Thus, measures of metabolism and activity in response to multiple stressors can provide additional insight into those stressors’ potential to affect other biological processes important to an individual’s fitness.

Although multiple stressors can interact in nearly all habitats, some such as the intertidal zone of rocky shores are characterized by especially unpredictable and extreme variation in multiple abiotic parameters, making them particularly useful model systems. In the intertidal zone these parameters include temperature, dissolved oxygen concentration, salinity, and pH (Denny and Dowd, 2022; Helmuth et al., 2002; Liguori, 2022; Morris and Taylor, 1983). Two of the most variable parameters in splashpools of the supratidal zone are salinity and temperature, which fluctuate over different timescales. Temperature oscillates over the day-night cycle, while salinity increases more gradually due to evaporation and can drop rapidly due to rainfall and wave splash (Denny and Dowd, 2022). On the west coast of North America, the copepod *Tigriopus californicus* inhabits isolated splashpools of the supratidal zone and has evolved to tolerate simultaneous variation in salinity and temperature. This copepod can tolerate salinities ranging from 2 to at least 190 ppt (Burton and Feldman, 1982; Denny and Dowd, 2022) and has been found in splash pools with temperatures higher than 40°C (Kelly et al., 2012), making it a useful model organism for studying the physiological effects of interactions between salinity and temperature.

Under an additive scenario, the expected pattern would be for survival and performance to decrease under simultaneous exposure to changes in salinity and high temperature. Changes in environmental salinity present a physiological challenge by disrupting ion gradients (Kültz, 2012; Somero et al., 2017), which imposes an energetic cost as the organism works to reestablish appropriate solute concentrations (Evans and Kültz, 2020; Somero et al., 2017; Thabet et al., 2017). Similarly, heat stress presents significant costs by disrupting biomolecular structures and increasing rates of biochemical reactions (Dahlhoff and Somero, 1993; Hoekstra and Montooth, 2013; Scheffler et al., 2019; Somero, 2020). When facing both of these challenges simultaneously, it would be expected under the additive framework that the costs of these stressors together would combine to decrease an organism’s aerobic scope, thus reducing its ability to maintain basal function, locomotion, growth, and reproduction (Sokolova, 2013).

*T. californicus* individuals exposed to simultaneous low salinities and high temperatures appear to respond similarly to additive expectations, where exposure to low salinity leads to a reduction in thermal tolerance (measured as lethal temperature for 50% of the sample, or LT_50_; Denny and Dowd, 2022; Kelly et al., 2016). However, contrary to predictions based on the energetic costs of responses to acute temperature or salinity increases separately (Goolish and Burton, 1989; Hoekstra and Montooth, 2013; Scheffler et al., 2019), *T. californicus* survival under simultaneous high temperature and salinity demonstrates a strong antagonistic interaction, where mortality is decreased relative to the dire anticipated additive scenario (Denny and Dowd, 2022). In the splashpool environment, where high salinities are often indicative of high temperatures in the recent past and greater potential for hot conditions in the near future, high salinities confer a level of cross-tolerance to high temperatures (Denny and Dowd, 2022). Similar patterns have been observed in other species within the genus, including *T. brevicornis* and *T. fulvus* (Damgaard and Davenport, 1994; Ranade, 1957). However, this interaction has yet to be directly investigated for any metric other than survival (but see Kelly et al., 2016).

Here, we describe the effects of salinity, temperature, and the interaction between them on the metabolic rate and activity of *T. californicus* by exposing copepods to changes in temperature under various acute and chronic salinity exposures. We hypothesized that responses in metabolic rate and activity to changes in temperature are non-additively influenced by acute changes in salinity levels, reflective of the cross-tolerance previously observed for survival. Based on the tenets of dynamic energy-budget theory, where allocation of energetic resources to one response decreases energy available for others (Kooijman, 2000), we predicted that increases in metabolic rate in response to low assay salinity would decrease resources available for addressing heat stress, explaining the low heat tolerance previously described at low salinities (Denny & Dowd, 2022). Conversely, we expected high assay salinities to be associated with lower metabolic rates, in line with results of other studies on *Tigriopus* species’ metabolic responses to salinity (McAllen and Taylor, 2001). We also expected that high assay salinities would interact antagonistically with high temperature, resulting in a smaller increase in metabolic rate relative to ambient or low assay salinities. Similar to predictions for metabolism, we anticipated that levels of activity would be negatively associated with temperature, as physiological and biochemical responses to increased temperatures draw on energy that would be otherwise allocated to activity. We predicted that high-salinity copepods would demonstrate higher levels of activity under high temperatures compared to low and ambient salinity copepods, due to the cross-tolerance between salinity and heat allowing for more energetic resources to be dedicated to activity. In partial support of our predictions, we found a strong impact of salinity, temperature, and their interaction on copepod activity, but less clear effects of temperature and its interaction with salinity on metabolic rates. Our findings suggest that activity of small copepods strongly influences metabolic rate and that changes in levels of activity may be an effective cost-saving strategy when faced with environmental stress.

## Methods

### Animal collection and culture maintenance

Laboratory cultures of *T. californicus* were established from a single collection at Cattle Point on San Juan Island in Washington state (48.452362°, –122.961969°). Copepods were collected on May 8^th^, 2023 from several independent splashpools and were transported to Washington State University in Pullman, WA. To create genetically homogenous groups across treatments, copepods from several splashpools were kept in their original seawater, mixed together, and subsequently separated into 16 separate 32 oz. jars (henceforth referred to as cultures). Cultures were kept at a constant 17.5 ± 0.5°C on a 12L:12D photoperiod. Over the course of several weeks, salinities were changed in cultures by 5 parts per thousand (ppt) twice a week until target treatment salinities were reached (20, 35, and 50 ppt, 4 cultures each; 65 and 80 ppt, 2 cultures each). The average salinity of seawater around Cattle Point is about 35 ppt. Although splashpools are not inundated with seawater during high tide, they do receive some water through wave spray that brings splashpools closer to this salinity; thus, 35 ppt groups serve as a control, representing the most typical natural conditions. After all cultures had reached target salinities (henceforth referred to as acclimation salinity), they were allowed to acclimate for at least two weeks to minimize effects from previous environmental history. Every two weeks, cultures within salinity treatments were mixed with one another to minimize genetic drift between cultures. Fresh water was added to cultures as needed to maintain salinities within 2 ppt of the given target salinity. Copepods were fed twice weekly *ad libitum* diets of ground algae pellets and TetraMin fish flakes. All experiments were conducted using only adult male copepods, which are distinguished by the lack of eggs and distinct morphology of the antennae (Vittor, 1971). Respirometry and heat-ramp activity experiments described below were completed between June 20^th^ and July 12^th^ of 2023, and acute salinity transfer activity experiments were completed between October 18^th^ and December 20^th^ of 2023, all using the same cultures. Since generation time of *T. californicus* is around 23-26 days (Edmands and Harrison, 2003), we assume that all experiments were on copepods in the F2 generation or later. Differences between F2 and later generations due to adaptation to the various acclimation salinities are unlikely, as previous work has shown no evidence for adaptation in this species to increased or decreased salinities over six generations (Kelly et al., 2016).

### Oxygen consumption measurement as a proxy for aerobic metabolic rate

A series of respirometry experiments were conducted to measure oxygen consumption upon chronic exposure to and acute transfer between salinities, as well as the effect of temperature on oxygen consumption at a range of salinities. Individual respirometry experiments were run using two Loligo® Systems 24-well glass microplates with 80µL wells and two PreSens SDR SensorDish® Readers. Before trials began, microplates were calibrated to each salinity and temperature combination using 0% and 100% oxygen saturation solutions, because the factory calibrations proved unreliable across this range of conditions. The 100% saturation saltwater solutions were made at their respective salinities by thoroughly shaking vials of solution, and 0% solutions were made by mixing saltwater at the respective salinity with sodium sulfite (1% w/v) and letting the solution absorb the oxygen for an hour. Columns of wells in each plate were filled with either the 100% or 0% oxygen solution such that there was an equal number of wells/columns with each solution. Plates were sealed and placed on the SDR reader within the incubator set to the respective temperature and measurements of oxygen saturation were taken every three minutes for thirty minutes. Phase readings from the SDR software (SDR_v4.0.0) for each well were then averaged across the 100% or 0% oxygen saturation groups and those results were used in the calibration settings for the respective salinity and temperature combination.

The day before a given respirometry trial, 14 copepods from each salinity were separated from cultures and placed in jars with artificial seawater at the same salinity as their culture, totaling 42 copepods per trial (for trials of chronic exposure to 65 and 80 ppt salinities, this was 23 copepods from each culture, totaling 46 per trial). Separated copepods were not fed for 24 hours before the experiment to avoid measurement of specific dynamic action (rates of oxygen in two calanoid copepod species return to baseline levels within 10 hours of feeding; Thor, 2000). On the day of a given trial, copepods were loaded into the microplates with one animal per well. Copepods were transferred with as little of their original water as possible into new jars of artificial seawater, filtered with a 0.22µm Millipore filter to minimize presence of algae, debris, or bacteria that would consume or produce oxygen independently of the copepods. Copepods were pipetted out of the filtered water along with 81µL of water and placed individually into their respective wells. One well of filtered water with no animal was included for each acclimation salinity on each plate to identify any background oxygen consumption. After all copepods had been loaded onto the microplates, a pipette tip was used to remove any air bubbles within wells. An adhesive microplate film with spots (made from inverted punches of the same film material) covering areas where the adhesive would otherwise come in contact with the water in the wells was then placed over each microplate to ensure an air-tight seal of each well. Each plate was then topped with a silicon sealing mat, a steel block, and a heavy plastic block to prevent any gas exchange during measurement. Plates were placed on a PreSens SDR reader inside an incubator set to either 17.5°C or 27.5°C. Dissolved oxygen (DO) content in each well was measured every three minutes over the course of 12 hours or until DO dropped below 80 Torr in every well. In preliminary trials, DO dropped below 80 Torr only in high temperature treatments and only after 7 or more hours. After a given respirometry experiment, individuals were taken from their wells, dabbed dry on a Kimwipe, and placed in a drying oven overnight. Dried individuals were then weighed to the nearest microgram to test for individual mass effects on calculated oxygen consumption rates. However, experiments were left to run overnight and some mortality during experiments under high-temperature treatments led to decomposition of those animals before dry mass could be measured. As a result, masses measured for the high temperature experiments may be less accurate. Metabolic rates from these animals were still used within the ranges of data described below, as within those ranges animals were still well above the critical P_O2_, discussed below.

Rates of oxygen consumption were calculated in R (v4.2.2; R Core Team, 2022) using the respR package (v2.3.1; Harianto et al., 2019) and the calc_rate() function, then subsequently converted to rates of ug hr^-1^ using the convert_rate() function. Data from the first 60 minutes of measurement were omitted to avoid measuring changes in metabolism due to handling and/or during temperature equilibration of the microplate. Data used were also limited to those collected within 10 hours of sealing the plate and when oxygen concentration in a well was above 30 Torr. This DO cutoff is well above the critical P_O2_ at which respiratory independence from oxygen concentration is lost for another species in the genus (∼16 Torr at 30°C and less at lower temperatures; McAllen et al., 1999). Thus, declines of DO in the wells above 30 Torr can be attributed to normal respiration, and we assume that DO content was not affecting oxygen consumption rate. This was further confirmed after analysis by a lack of a consistent temporal effect on metabolic rates across treatments. Rates of each acclimation salinity treatment within a plate were corrected by subtracting the background consumption rates of the blank wells with water from the corresponding acclimation salinity. Individual rates were plotted against masses in the low temperature treatments, and there appeared to be no allometric relationship between the two variables (Fig. S1). Because of this and the potential for inaccuracy in the measured masses of high temperature animals, we forwent any mass-correction of measured rates.

### Oxygen consumption upon acute salinity transfer

We measured changes in routine metabolic rates (RMR) in response to acute salinity changes under ambient (17.5°C) or high (27.5°C) temperatures to disentangle metabolic costs due to maintenance under various salinities (chronic salinity exposure) and the cost of adjusting solute concentrations in response to an acute change in salinity. Copepods from 20, 35, and 50 ppt cultures were transferred from their acclimation salinities to a common assay salinity (20, 35, or 50 ppt) immediately before being loaded into the microplate (Fig. 1A). In these trials, copepods were loaded into wells in respirometry plates starting with those already acclimated to the target salinity (e.g., if an assay salinity was 20 ppt, copepods from 20 ppt cultures were loaded first) so as to minimize the time that acute transfer copepods (e.g., 35 or 50 ppt acclimation copepods in 20 ppt assay salinities) had to adjust to the new salinity before measurements could begin. Assay salinity for a given trial was constant across wells and plates, and placement of copepods from the three acclimation salinities was randomized across the plate.

**Figure 1.**
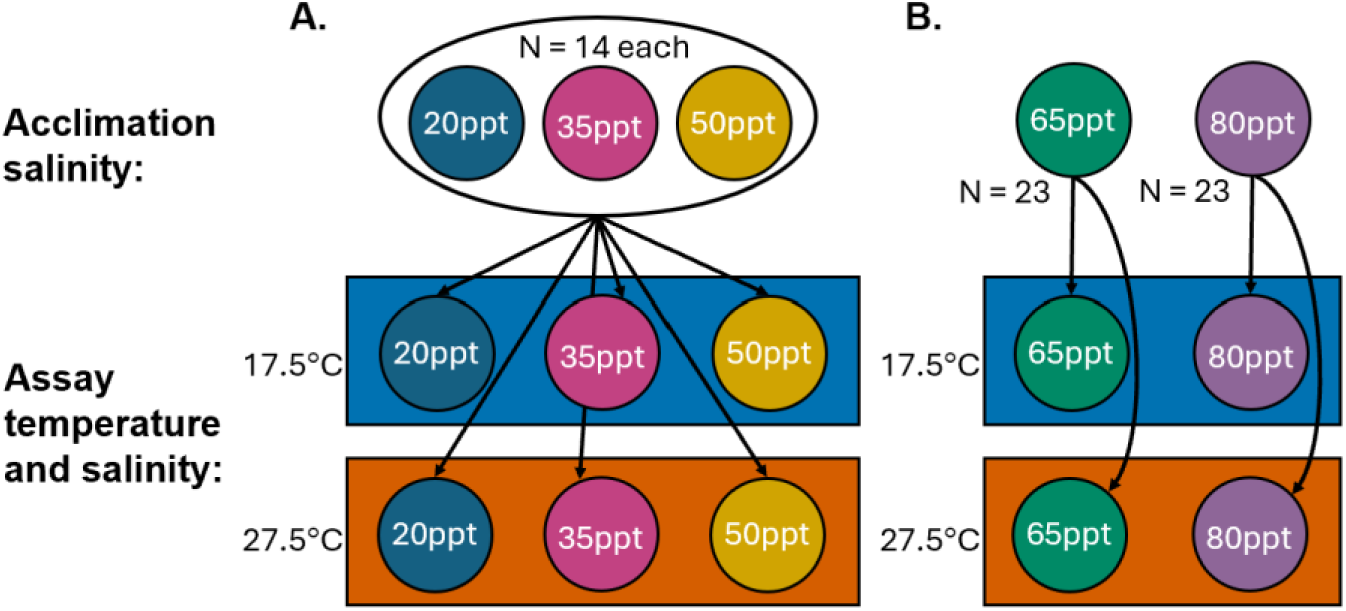
Respirometry experimental design. **A)** Acute salinity transfer, where in each of two microplates, 7 copepods from each acclimation salinity were transferred to a common assay temperature and salinity. **B)** High salinity treatments where no acute transfers between salinities were performed, and two high acclimation salinities were assayed in two temperatures.

### Oxygen consumption at high salinities

Although *T. californicus* can tolerate extremely high salinities and have been found in splashpools with salinities as high as 190 ppt (Denny and Dowd, 2022), effects of salinity on metabolic rate in this species have only been measured in salinities as high as 60 ppt (DeBiasse et al., 2018). Thus, in addition to measurement of routine metabolic rates of 20, 35, and 50 ppt copepods upon transfer to different salinities, we also measured rates of oxygen consumption of copepods chronically acclimated to higher salinities (65 and 80 ppt) in the same ambient and high temperatures (Fig. 1B). More southern populations of this species can survive up to 190 ppt, but our northern Washington field site salinities do not reach such extremes. As such, 80 ppt is near the limits of their chronic salt tolerance (C. Terry, unpublished observations). Effects of acute transfers to/from these salinities were not measured, because changes of such a large magnitude would be unrealistic in nature and very stressful, if not lethal. Indeed, in previous work unrealistic acute transfers between salinities (e.g., a 70 ppt shift from 30 to 100 ppt) have not resulted in the increase in thermal tolerance that is seen under more natural salinity changes (e.g., a 30 ppt shift from 30 to 60 ppt; Denny & Dowd, 2022). In these trials, copepods were prepared as described above and transferred to filtered water of the same salinity before being placed in their individual wells on the microplate. Measurements of DO content were taken in the same manner as described above, except measurements for 65 and 80 ppt copepods were collected at the same time, with each salinity allotted to its own plate.

### Analyses of oxygen consumption data

After all experiments, rates of individual oxygen consumption in µg hr^-1^ were fitted to linear mixed effects models using the lme4 package in R (Bates et al., 2015). To evaluate the effects of chronic salinity acclimation and temperature on copepod metabolic rates, we combined data from chronically acclimated high-salinity copepods (65 and 80 ppt) with same-salinity transfer individuals (i.e., no acute salinity changes). These chronic acclimation data were fitted to a model with fixed terms for acclimation salinity and temperature and a random effect for microplate. Significance of model terms was determined using type-three sums of squares in the car package (v3.1-2; Fox et al., 2023). Significant differences between groups within a treatment were compared using pairwise comparisons with a Tukey adjustment in the emmeans package (Searle et al., 1980). To better understand the effects of acute temperature changes, another linear mixed effects model was fitted with the same oxygen consumption data, but with RMR calculated over hour-long intervals as opposed to over the entire 9-hour period. Data were fitted to a similar model as the overall data, but with an added fixed effect of hour and an added random effect of individual ID to account for repeated measures.

To evaluate the effects of acute salinity transfer under different temperatures, data from all copepods in the acute transfer experiments (i.e., no 65 and 80 ppt copepods) were fitted to a model with fixed terms for acclimation salinity, assay salinity, and temperature and a random effect for microplate. Significance of model terms was determined using type-three sums of squares, and significant (p-value < 0.05) two-way interactions were further evaluated using pairwise comparisons with a Tukey adjustment. Similar to chronic acclimation copepods, the effects of acute salinity changes were further investigated by fitting another model with RMRs calculated over hour-long intervals. This model included an added fixed effect of hour and an added random effect of individual ID to account for repeated measures.

### Measurement of rates of activity

Activity levels of copepods were measured using a novel application of a multimode microplate reader (Tecan Spark; Tecan Trading Ltd., Switzerland) developed previously in our lab (Cuevas-Sanchez, 2019). The instrument measures absorbance of a narrow beam of infrared light near the edge of each well of a 96-well plate. *T. californicus* is unable to perceive light in the infrared spectrum, and previous lab observations have shown that when placed in a well, these benthic copepods largely spend their time making laps around the edges of the wells. Thus, instances of infrared light absorbance can be interpreted as a copepod’s completion of a lap around a well, and we assume that the light has no effect on their activity. The instrument is also able to control internal temperatures and carry out measurements under a ramping temperature regime, making it ideal to measure changes in activity across temperatures. In these activity assays, copepods along with 200 µL of water were placed randomly into individual wells of a 96-well plate, covered with a transparent adhesive film to prevent evaporation, and placed in the instrument at 18°C. After 30 minutes under constant conditions, measurements at 18°C began. In both assays copepod activity was recorded for three minutes, three times per individual at each step (each temperature or time block), cycling through the individuals to avoid systematic bias due to factors such as time at temperature. After each assay, survival was documented and only data from surviving copepods were used for analysis. To obtain activity counts from the absorbance peaks, data were run through a custom MATLAB script using the findpeaks() function in the Signal Processing Toolbox (The MathWorks Inc., 2017). The peak detection parameters were experimentally tuned: MinPeakDistance was set to 1, MinPeakHeight was set to 0.01, and MinPeakProminence was set to 0.007. Peak identification was visually checked for all data, and individuals’ counts of activity (in number of peaks) were summed across the three three-minute measurement intervals at each step (temperature or time block).

### Activity under increasing heat

To quantify changes in activity levels of *T. californicus* in response to increased heat under various chronic salinity treatments, we conducted activity assays using only those adult male copepods chronically acclimated to 20, 35, 50, 65, and 80 ppt. This assay allowed for investigation of the presence of any interaction between temperature and acclimation salinity in activity responses. Spread over the course of seven assays, a total of 21 copepods from each salinity underwent a step heat ramp (Fig. 2). Each heat ramp assay included three individuals from each salinity. Measurements began at 18°C and continued over the course of five temperature steps total, ending after measurement at 30°C. Between steps, temperature was increased by 3°C over the course of three minutes and held at the target temperature for 15 minutes before measurements at that step began. After the last measurement at 30°C, temperature was returned to 18°C until copepods were checked for survival. Activity counts from MATLAB were fitted to a generalized linear mixed effects model in R using the lme4 package, with fixed effects of temperature, salinity, and their interaction, a random effect of individual ID to account for repeated measures, and a Poisson link function due to the use of count data. Predicted counts of activity for each salinity at each temperature were calculated using the model and the ggpredict() function in the ggeffects package (v1.52; Lüdecke, 2018). Significance of model terms was determined using type-three sums of squares.

**Figure 2.**
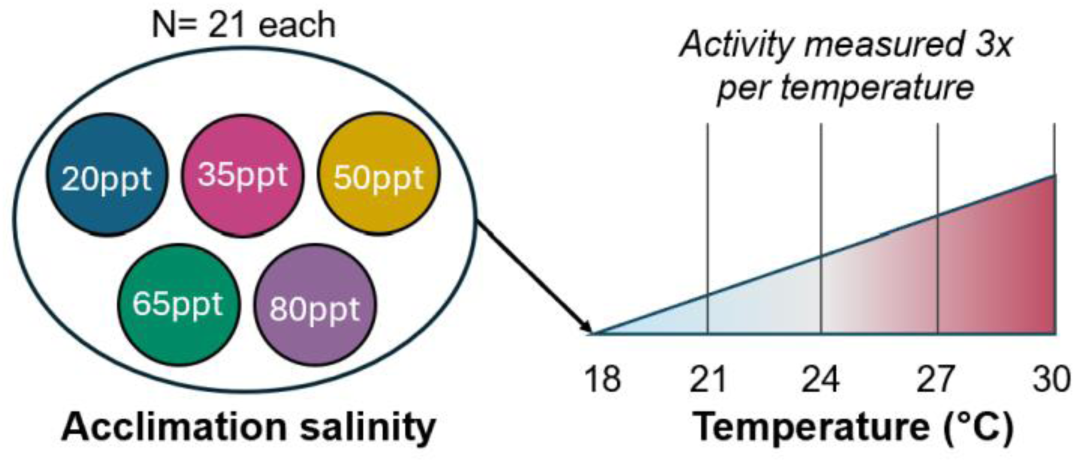
Activity under increasing heat experimental design. In each trial four copepods from each acclimation salinity underwent a step heat ramp from 18 to 30°C. Activity was measured three times at each temperature.

### Activity after acute salinity transfer

To capture any effects of acute salinity transfer and subsequent short-term responses on copepod activity, we measured activity rates of *T. californicus* upon acute transfer among 20, 35, and 50 ppt. This was tested with a fully factorial design similar to that of the previously described respirometry assays (Fig. 3). Five copepods from each acclimation salinity were individually transferred to microplate wells filled with 200µL of a common assay salinity. Activity was measured at the same time intervals as the heat ramp assays, but without increasing temperatures (i.e., at a constant 18°C throughout). Thus, measurement times were grouped into 5 corresponding time intervals: 0.5-2.75 hours, 3-5.25 hours, 5.5-7.75 hours, 8-10.25 hours, and 10.5-12.75 hours. Four assays were completed at each of the three assay salinities, resulting in 20 individuals in each crossed-treatment combination. Resulting activity counts were fitted to a generalized linear mixed effects model using the lme4 package with fixed effects of time, acclimation salinity, assay salinity, and their interactions, a random effect of individual ID to account for repeated measures, and a Poisson link function. Predicted counts of activity for each salinity at each time point were calculated using our model and the ggpredict() function. Significance of model terms was determined using type-three sums of squares.

**Figure 3.**
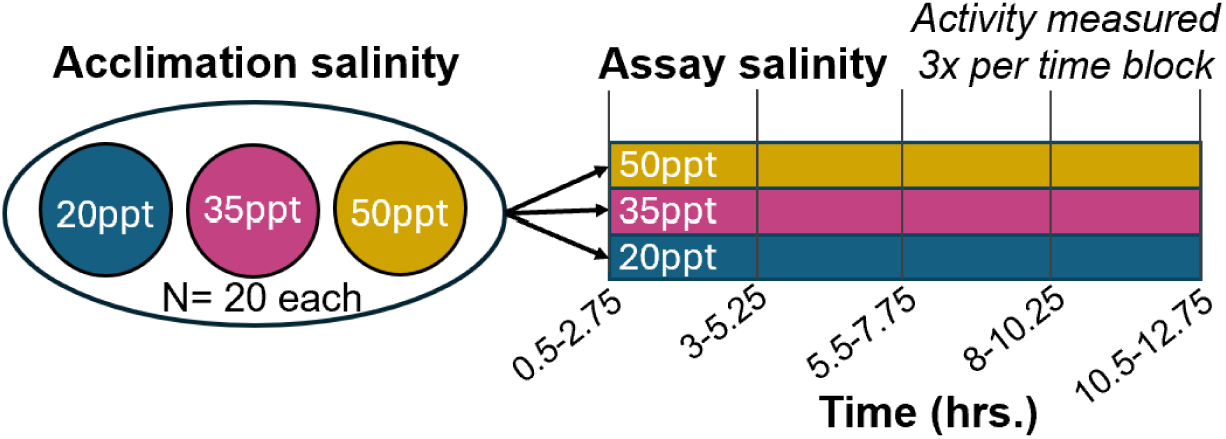
Experimental design for measuring activity after acute salinity transfer. In each trial five copepods from each acclimation salinity were acutely transferred to a common assay salinity and activity was measured at the same time points as in the heat ramp assays.

## Results

### Chronic increases in salinity reduced copepod metabolic rates, but temperature change had no effect

Routine metabolic rates (RMR) of chronically acclimated copepods in all five salinity treatment groups were measured at both ambient and high temperatures. When considering metabolic rates calculated over the entire experimental period, salinity strongly influenced RMR (p < 0.0001; Table 1), demonstrated by an inverse relationship between salinity and RMR at both temperatures (Fig. 4). Between 20 and 80 ppt treatments, mean RMR decreased by 23% and 28% in 17.5°C and 27.5°C, respectively. In contrast with our expectations, temperature had no effect on RMR, and there was no interaction between salinity and temperature (Table 1). Post-hoc pairwise comparisons averaged over temperature levels resulted in significant differences between all salinity treatments except for the 35 and 50 ppt comparison (Fig. 4; Tukey pairwise comparison: p = 0.929). In the hour-by-hour analyses, there was no visually obvious initial increase in metabolic rates in chronically acclimated copepods at either temperature (Fig. S2). This indicated that there was no increased respiration from handling stress or exposure to the acute temperature increase in the period measured (beginning an hour after placement in the incubator). Nonetheless, there was a statistically significant effect of hour (p = 0.0187; Table S1) and an interaction between hour and temperature (p = 0.0012; Table S1), but these effects were small and not consistently apparent in the beginning of the measurement period where acute temperature effects would be expected (Fig. S2).

**Figure 4.**
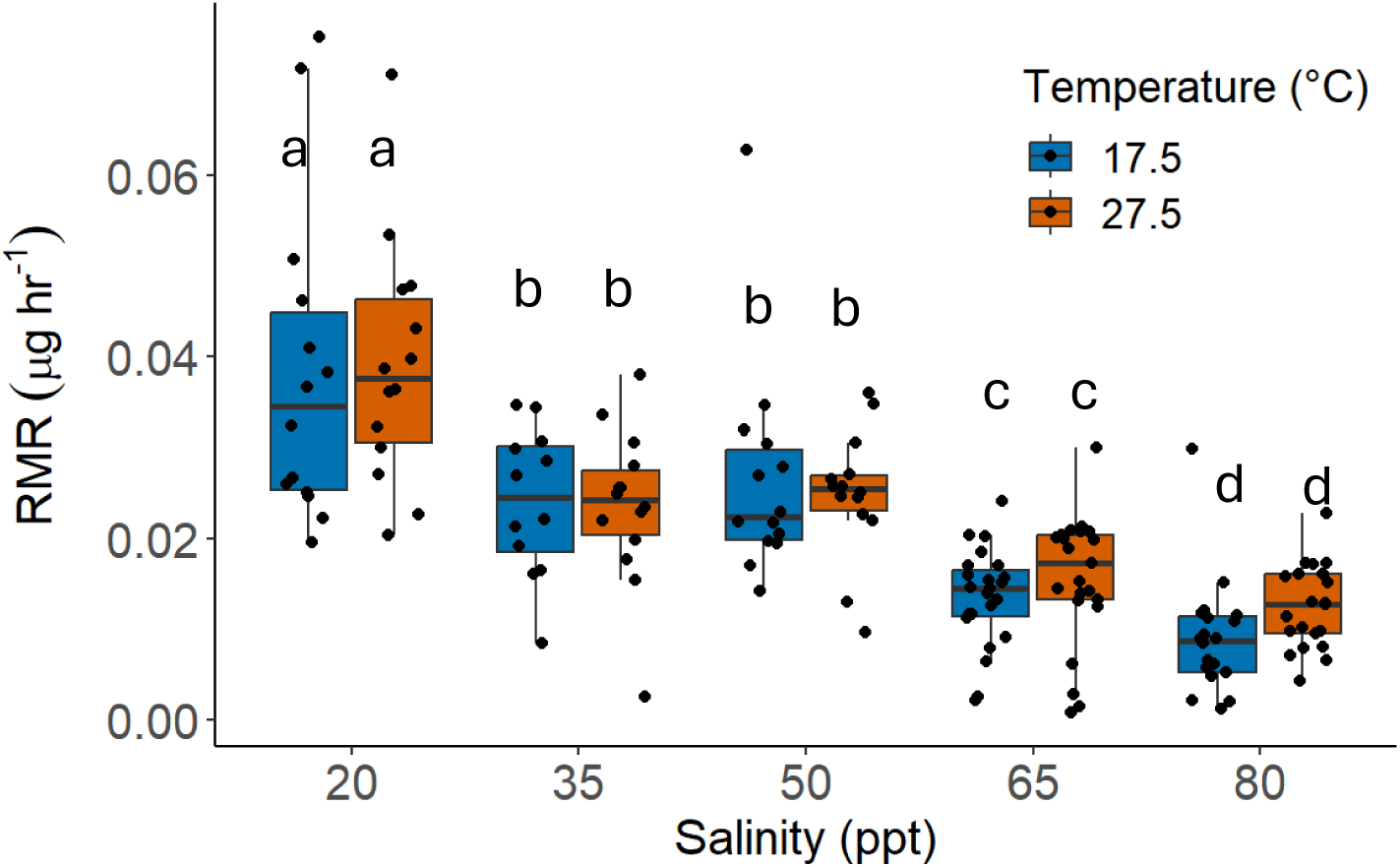
Routine metabolic rates (RMR) of adult male copepods chronically acclimated to their respective salinities and measured in ambient (17.5°C) and high (27.5°C) temperatures. Median values are depicted by black lines in the middle of boxes, upper and lower box hinges correspond to the first and third quartiles, and whiskers extend to no further than 1.5 times the interquartile range. Letters indicate groups significantly differing in RMR, as determined by Tukey pairwise comparisons.

**Table 1.** Summary of the statistical linear mixed effects model results testing for effects of chronic salinity acclimation, temperature, and their interaction on copepod RMR.

| Factor | Chi square value | Df | p-value |
| --- | --- | --- | --- |
| **Salinity** | **125.0057** | **4** | **< 2e-16** |
| Temperature | 0.2957 | 1 | 0.5866 |
| Salinity*Temperature | 1.4187 | 4 | 0.8409 |

Significant terms are bolded.

### Acute transfers to lower salinities initially increase copepod metabolic rates

Acute transfer of copepods acclimated to 20, 35, and 50 ppt among each of those salinities had little effect on RMR when calculating rates of oxygen consumption over the entire measurement period. Acclimation salinity and assay salinity both had significant main effects on RMR (Table 2). There was a significant interaction between acclimation salinity and assay salinity (p < 0.0001; Table 2), but subsequent one-way ANOVAs and post hoc Tukey comparisons within assay salinities suggest that this interaction is driven solely by copepods acclimated to 50 ppt and transferred to 20 ppt (Fig. 5). Specifically, within assay salinities there was a significant effect of acclimation salinity only within the 20 ppt assay groups (p < 0.0001 at 17.5°C, p = 0.0072 at 27.5°C), and post hoc Tukey comparisons demonstrated that RMRs of copepods acclimated to 50 ppt were significantly higher than the other two acclimation salinities at both temperatures (Fig. 5). Interestingly, acute transfer in the reverse direction from 20 to 50 ppt had no influence on RMR. Like chronically acclimated copepods, there was no main effect of temperature on RMR after acute salinity transfer, but there was a significant two-way interaction between acclimation salinity and temperature and a significant three-way interaction (acclimation salinity*assay salinity*temperature; Table 2). When RMR was analyzed on an hourly basis, increased metabolic costs due to acute salinity transfer became apparent, but only in transfers from higher to lower salinities (Fig. 6). Increases in RMR after acute transfer were greatest in transfers of the greatest magnitude (i.e., 50 to 20 ppt) and in transfers to 20 ppt. The time until metabolic recovery also seemed to follow this trend, such that RMRs of copepods in the 50 to 35 ppt transfer group appeared to plateau earlier than those in the other two groups (Fig. 6). When comparing outcomes of the model of overall rates (Table 2) to the model that used rates calculated over hour-long intervals (Table S2), there was a large decrease in the amount of variance accounted for by the three-way interaction between assay salinity, acclimation salinity, and temperature, such that it was no longer significant (p = 0.1846; Table S2). The main effect of assay salinity followed the same trend between the models and remained marginally significant in the hour-by-hour model (p = 0.0557; Table S2). In the model with hour-by-hour measurements of RMR, there were also additional significant effects from two-way interactions between acclimation salinity and hour (p < 0.0001; Table S2) and temperature and hour (p = 0.0224; Table S2), and a significant three-way interaction between acclimation salinity, assay salinity, and hour (p = 0.0003; Table S2).

**Figure 5.**
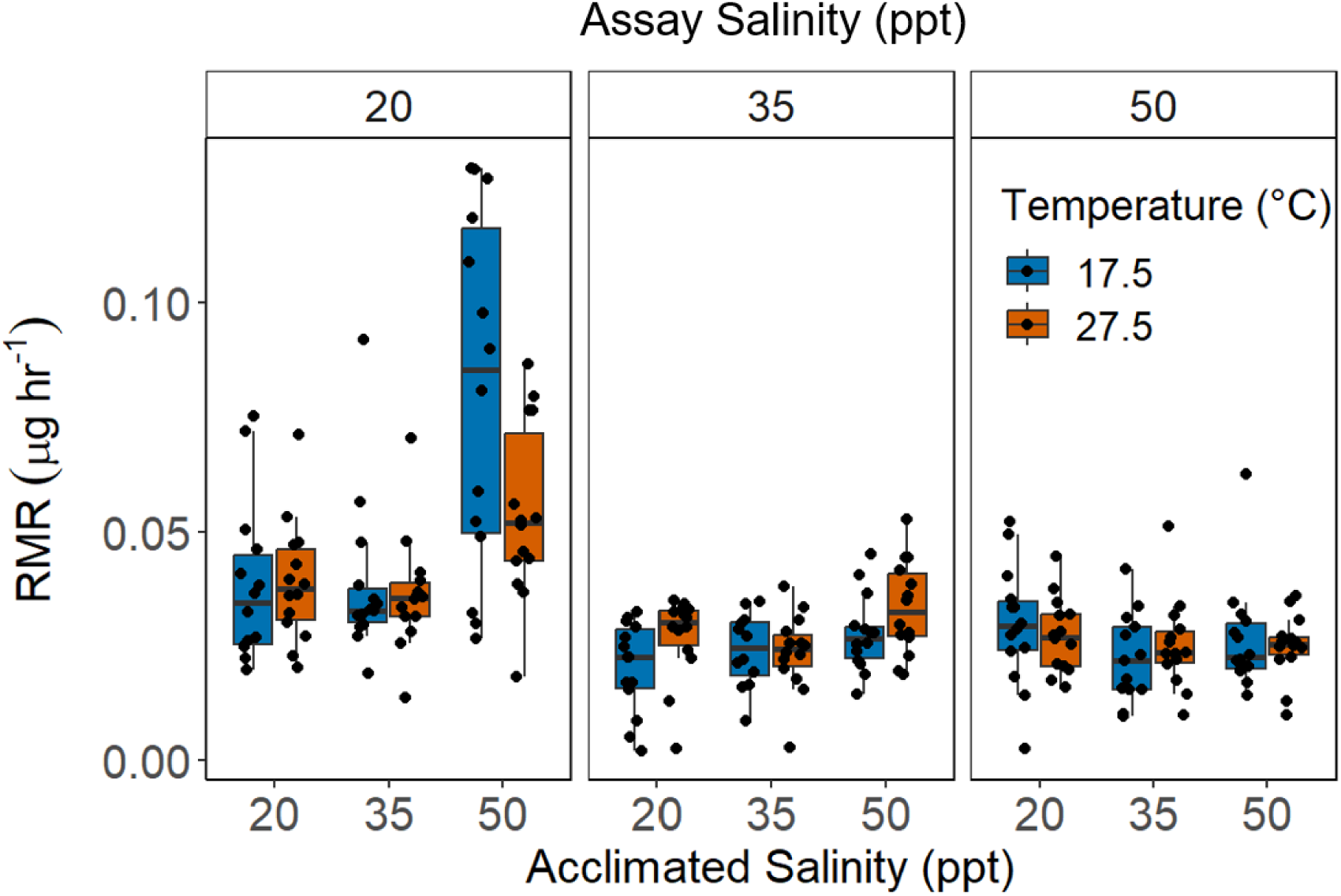
Routine metabolic rates (RMR) of chronically acclimated, adult male copepods that were acutely transferred between salinities and measured in both ambient and high temperatures. Facets represent a given assay salinity. Median values are depicted by black lines in the middle of boxes, upper and lower box hinges correspond to the first and third quartiles, and whiskers extend to no further than 1.5 times the interquartile range.

**Figure 6.**
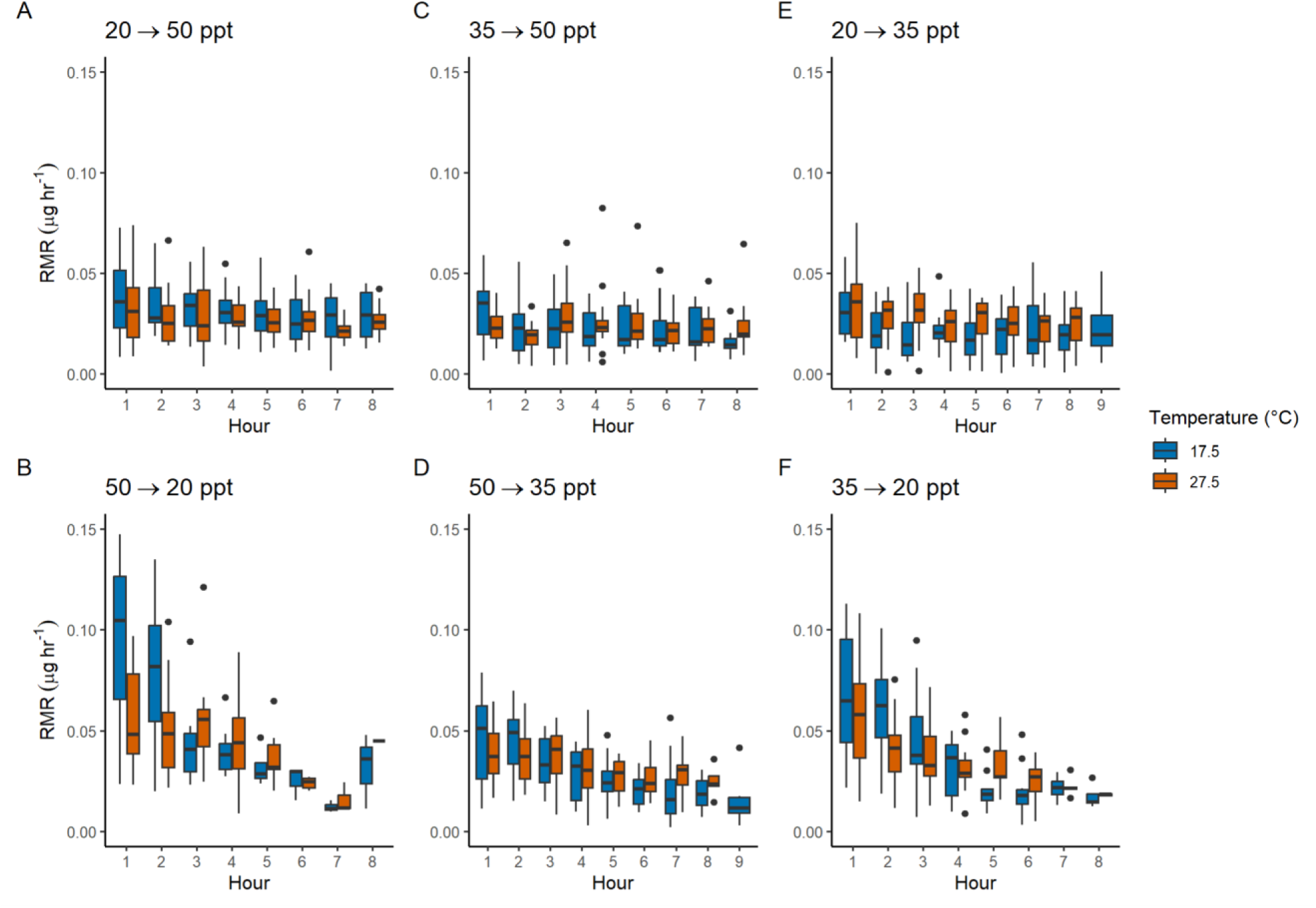
Hourly measures of routine metabolic rates (RMR) of chronically acclimated, adult male copepods that were acutely transferred between salinities and measured in both ambient and high temperatures. Panels represent a given salinity transfer (acclimation salinity → assay salinity). Median values are depicted by black lines in the middle of boxes, upper and lower box hinges correspond to the first and third quartiles, and whiskers extend to no further than 1.5 times the interquartile range. Metabolic rate rose initially upon salinity decrease (B, D, F), but not upon salinity increase (A, C, E).

**Table 2.** Summary of the statistical linear mixed effects model results testing for effects of acute salinity transfer and temperature on copepod RMR.

| Factor | Chi square value | Df | p-value |
| --- | --- | --- | --- |
| **Acclimation salinity** | **79.9422** | **2** | **< 2e-16** |
| **Assay salinity** | **8.1795** | **2** | **0.0167** |
| Temperature | 0.1062 | 1 | 0.7445 |
| **Acclimation salinity\*Assay salinity** | **47.0819** | **4** | **1.466e-09** |
| **Acclimation salinity\*Temperature** | **14.3329** | **2** | **0.0008** |
| Assay salinity*Temperature | 0.7808 | 2 | 0.6768 |
| **Acclimation salinity\*Assay salinity\*Temperature** | **10.6216** | **4** | **0.0312** |

Significant terms are bolded. Analysis excludes copepods acclimated to 65 and 80 ppt and includes copepods chronically acclimated to 20, 35, and 50 ppt and then acutely transferred to each of those three salinities.

### Increasing temperatures and salinities decrease copepod levels of activity

Copepods chronically acclimated to all five salinities were exposed to a heat ramp from 18°C to 30°C over the course of about 12 hours. Rates of activity showed a strong negative response to both increasing salinities and increasing temperatures (p < 0.0001 for both salinity and temperature; Table 3). Activity levels of copepods acclimated to 80 ppt were at most 40% of those in the lowest salinity (20 ppt) across all temperatures (Fig. 7). For copepods acclimated to 20 ppt, predicted levels of activity in the model dropped by 52.6% between the lowest and highest temperatures. Across the five salinities, this decrease in activity ranged from 46.5% in 35 ppt copepods to 87.8% in 65 ppt copepods (Fig. 7). Temperature and acclimation salinity also significantly interacted to affect activity (p < 0.0001; Table 3). This difference in temperature responses between salinities is most apparent in copepods acclimated to 35 ppt, for which activity changed the least across temperatures (Fig. 7).

**Figure 7.**
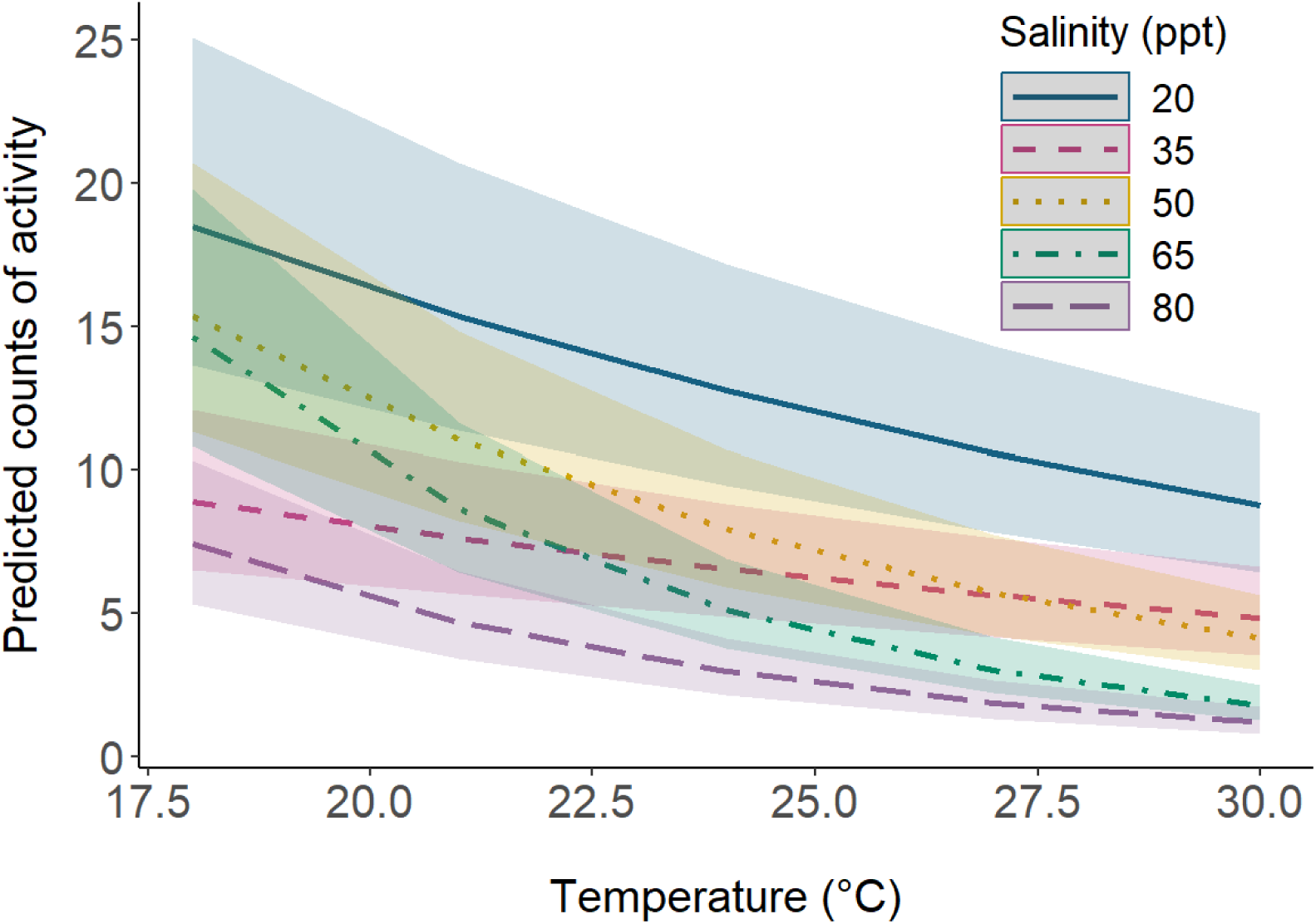
Copepod activity declines with increasing salinity and temperature. Predicted counts of activity are from a generalized linear mixed model with fixed effects of salinity, temperature, and an interaction between the two, and a random effect of individual. Colored ribbon around each line denotes the confidence interval for predictions at that salinity and temperature. All raw data are presented in Fig. S3.

**Table 3.** Summary of the statistical linear mixed effects model (with individual as a random factor) results testing for effects of chronic salinity acclimation, temperature, and their interaction on copepod activity.

| Factor | Chi square value | Df | p-value |
| --- | --- | --- | --- |
| **Salinity** | **34.148** | **4** | **6.948e-07** |
| **Temperature** | **685.965** | **1** | **< 2.2e-16** |
| **Salinity\*Temperature** | **154.400** | **4** | **< 2.2e-16** |

Significant terms are bolded.

### Acute salinity transfer has no effect on copepod levels of activity

Acute transfers among 20, 35, and 50 ppt salinities appeared to have little effect on copepod activity levels. Both time after transfer and acclimation salinity had no significant effect on rates of activity, although there was a significant interaction between the two factors (p = 0.0018; Table 3). Instead, assay salinity and the interaction between assay salinity and time after transfer appear to have the most notable impacts on activity (Fig. 8; Table 3). Additionally, there was a significant three-way interaction between time after transfer, acclimation salinity, and assay salinity (p = 0.0001; Table 3).

**Figure 8.**
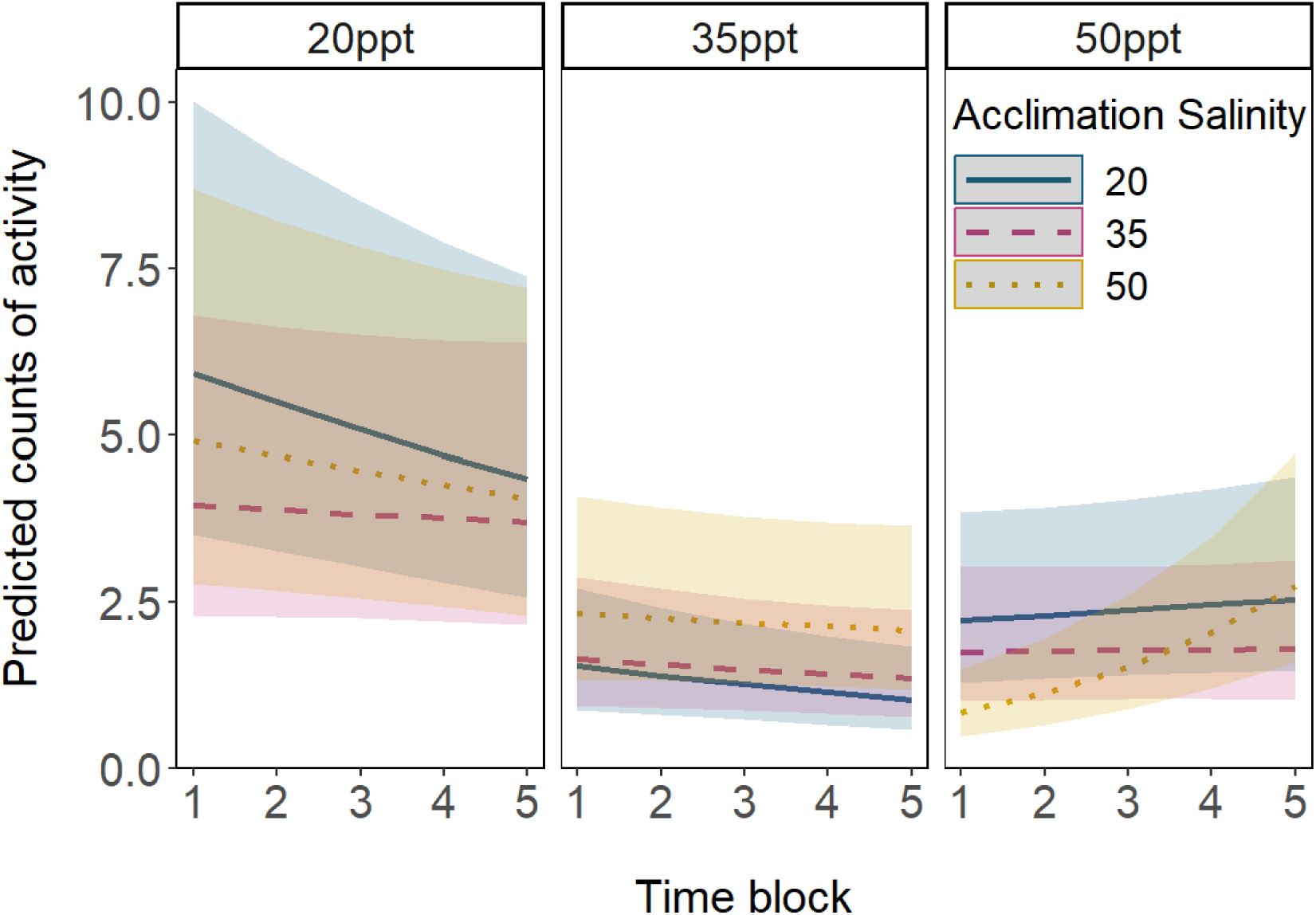
Acute salinity transfer impacts rates of copepod activity. Predicted counts of activity are from a generalized linear mixed model with fixed effects of acclimation salinity, assay salinity, time after transfer, interactions between all fixed effects, and a random effect of individual. Colored ribbon around each line denotes the confidence interval for predictions at that salinity and temperature. Assay salinities are represented by plot facets. All raw data are presented in Fig. S4.

## Discussion

The ephemeral splashpools that *Tigriopus californicus* inhabits in the supratidal zone experience drastic co-variation in salinity and temperature over a range of timescales (Denny and Dowd, 2022). These abiotic factors interact non-additively to affect acute heat tolerance, but the existence of such interactive effects on sublethal performance responses has not been previously reported. Here, we found that increased salinity decreased rates of both activity and oxygen consumption in *T. californicus*. Elevated temperature led to reduced rates of activity overall, resulting in no discernable impact of increased temperature on routine metabolic rates. Temperature also interacted with assay salinity to affect activity and with acclimation salinity to affect routine metabolic rates upon acute salinity transfer, implying that the sublethal impacts of these co-varying factors are also not predictable from experiments that study them in isolation.

### Effects of temperature on activity and routine metabolic rate

One of the notable and surprising findings of this study is the absence of temperature main effects on routine metabolic rates. With the 10°C increase in temperature between treatments in our respirometry experiments, it would be expected that rates of oxygen consumption would increase due to the Arrhenius relationship, where increased temperature increases rates of biochemical reactions (Somero et al., 2017). This phenomenon is also known as the Q_10_ effect, or the fold change in reaction rates with a 10°C increase in temperature. It is assumed that most ectotherms demonstrate Q_10_ values of 2-3 (Scholander et al., 1953), making *T. californicus’* lack of metabolic response to temperature surprising. However, some eurythermal ectotherms may have a lower Q_10_ of 1-2 (Branch et al., 1988; Scholander et al., 1953). Given the highly eurythermal nature of *T. californicus*, a low Q_10_, although surprising, is not entirely unexpected.

Current literature on the *Tigriopus* genus’ temperature sensitivity in metabolic rates provides conflicting evidence. Routine metabolic rates of *T. brevicornis*, a similarly eurytolerant congener that inhabits coastlines of northern Europe and survives salinities of 5 to 150 ppt and temperatures from –1 to 32°C, have been shown to exhibit an average Q_10_ value of 2.9 (McAllen et al., 1999). Within *T. californicus*, isolated mitochondrial rates of ATP synthesis and state 4 mitochondrial respiration rates increase with temperature across populations in southern California (Harada et al., 2019). These observations suggest notable temperature sensitivity in both whole-organism and mitochondrial respiration rates in the genus but are not in line with our current findings or those of some other work in *T. californicus*. In a study of metabolic responses to temperature change, Scheffler et al. (2019) found that copepods chronically exposed to a range of temperatures demonstrated no differences in metabolic rates (Q_10_ ≤ 1), but rates differed after acute increases in temperature. However, this acute temperature response was not present in one of the two northern populations in their study, and metabolic rates recovered within two to four hours of heat exposure in the other two populations. To determine whether there were more rapid changes in RMR upon acute temperature increase that would otherwise be undetectable over a 9-hour measurement period, we reexamined our oxygen consumption data in hour-long intervals. In line with the one northern population in Scheffler et al.’s study, we saw no notable differences in the first few hours of oxygen consumption between copepods exposed to different temperatures within a given chronic salinity acclimation group.

Scheffler et al. (2019) proposed that this rapid metabolic compensation to high temperatures in *T. californicus* may be due to fast-acting biochemical responses, but results of the current study suggest that a substantial amount of this effect may instead be driven by changes in activity. This species of copepod is very small (1-2 mm) and continuously active, and it is consequently impossible to measure standard metabolic rates – rates measured when no locomotor activity is occurring – without the use of drugs or restraints that might also impact physiology. Thus, any measurement of whole-organismal respiration rates in these copepods is subject to the effect of varying levels of activity (including those in the studies described above). The energetic cost of locomotory activity has been experimentally shown to comprise a large proportion of RMR in a range of aquatic species, including crucian carp and North Pacific krill (Nilsson et al., 1993; Torres and Childress, 1983). A model of calanoid copepod (6 mm length) swimming behavior estimated the cost of maximum velocity locomotion to be over 3 times that of standard metabolic rates (Morris et al., 1985). Given this evidence for non-negligible costs of locomotion in copepods and the strong decrease in activity with increases in temperature across salinities in our study (in most salinities, activity rates were more than halved between 18 and 30°C; Figure 3), it is likely that decreases in the energetic costs of activity are offsetting any increase in metabolic rates that would otherwise present as Q_10_ effects. These reduced rates of activity in high temperatures and high salinities may even suggest a cost-saving strategy in this species under stressful conditions. A similar strategy of decreased locomotor activity is utilized by overwintering fish in temperate regions to reduce metabolic costs, as an alternative to metabolic rate depression (Speers-Roesch et al., 2018). In *T. californicus,* reducing costs of locomotion in high salinities and temperatures may conserve energy for mounting biochemical responses to these stressors, perhaps contributing to this copepod’s increased survival under simultaneous high temperature and salinity (Denny and Dowd, 2022). However, ecological and other consequences of this behavioral adjustment remain to be determined.

### Effects of chronic salinity acclimation and acute salinity change on activity and routine metabolic rate

In respirometry experiments of copepods chronically exposed to a range of salinities, we found a strong decrease in metabolic rates with increasing salinity. Like decreased activity under high temperatures possibly offsetting Q_10_ effects in metabolic rates, decreased activity in high salinities may account for some of the decreases in metabolic rates. Similar trends in metabolic rates and activity under varying acclimation salinities are seen in *T. brevicornis*, such that metabolic rates and swimming behaviors are higher in low salinities and lowest in high salinities (McAllen and Taylor, 2001). McAllen and Taylor (2001) proposed that this decrease in activity in higher salinities may be due to higher viscosity of the medium and increased body density due to increased ion content. They posit that increased costs of locomotion under these conditions limit activity and thus result in lower metabolic rates. However, effects of salinity on water viscosity are outweighed by effects of increased temperature on viscosity (Chen et al., 1973), and there are no current experimental studies investigating effects of water viscosity on behavior and respiration of small marine organisms.

In contrast with chronic salinity exposures, acute salinity transfers for the most part had no effect on overall routine metabolic rate measured over several hours, with the exception of copepods acclimated to 50 ppt and transferred to 20 ppt. This result also contradicts findings for the congener *T. brevicornis*, for which there were significant responses in rates of oxygen consumption to acute transfers between salinities (McAllen and Taylor, 2001). This result is surprising, given the expected osmoregulatory costs associated with compensating for shifting ion and water movements at different salinities. For example, *T. californicus* exposed to changes in salinities alter internal concentrations of free amino acids over the following 24 hours (Burton and Feldman, 1982). Accumulation of proline and alanine, the two major osmolytes that are regulated in response to salinity in this species, in response to hyperosmotic stress is predicted to account for about 23.4% of daily energy expenditure in *T. californicus* (Goolish and Burton, 1989). Like metabolic responses to temperature, the temporal duration of our overall RMR experiments may have obscured the cost of osmolyte accumulation, as free amino acid concentrations change most rapidly within the first 6 hours of hyperosmotic exposure (Burton and Feldman, 1982). Thus, to investigate the time course of metabolic responses to acute salinity change, we reanalyzed these post-transfer RMR data over hour-long intervals. Interestingly, there were no initial increases in RMRs of copepods transferred to higher salinities, indicating either that this cost of acclimation to higher salinities is small, or that decreases in activity upon acute salinity increases obscure evidence of increased metabolism due to amino acid synthesis. This may explain the small and delayed upward trend of activity rates over time in copepods transferred to 50 ppt (Fig. 8) and the overall interaction seen between assay salinity and time (Table 4).

**Table 4.** Summary of the statistical linear mixed effects model (with individual as a random factor) results testing for effects of acute salinity transfer over time on copepod activity. Significant terms are bolded.

| Factor | Chi square value | Df | p-value |
| --- | --- | --- | --- |
| Acclimation salinity | 0.6746 | 2 | 0.7137 |
| **Assay salinity** | **23.3419** | **2** | **8.538e-06** |
| Time after transfer | 0.8802 | 1 | 0.3482 |
| Acclimation salinity*Assay salinity | 3.1418 | 4 | 0.5344 |
| **Acclimation salinity\*Time after transfer** | **12.6764** | **2** | **0.0018** |
| **Assay salinity\*Time after transfer** | **35.5084** | **2** | **1.947e-08** |
| **Acclimation salinity\*Assay salinity\*Time after transfer** | **22.8832** | **4** | **0.0001** |

In the case of hypoosmotic transfers or challenges, the cost of degrading free amino acids and maintaining intracellular ionic balance in response to low salinities may be greater than the cost of accumulating them. This may explain why the only notable acute transfer effect in this experiment (when considering RMR over the entire experimental period) was in the copepods transferred from 50 ppt to 20 ppt, the most extreme high-to-low transfer. Although, to the best of our knowledge, costs of free amino acid degradation in this genus have not been investigated, evidence in other crustacean systems suggests that there may be notable costs associated with amino acid catabolism during acclimation to decreases in salinity. Gilles (1973) showed that isolated axons of blue crabs decreased oxygen consumption upon hyperosmotic exposures and increased oxygen consumption upon hypoosmotic exposure; the latter change was attributed to increased rates of amino acid catabolism that correlated with increased oxygen consumption. A metabolomics-based study on another decapod crustacean, *Scylla paramamosain*, demonstrated a strong enrichment of amino acid metabolism pathways in response to acute drops in salinity (Yao et al., 2020). In an opposite pattern to the observations in *Tigriopus* species, reduced rates of respiration and activity under low salinities have been demonstrated in the intertidal flatworm *Macrostomum lignano* (Rivera-Ingraham et al., 2016). In that study, acute transfer to hypoosmotic conditions decreased activity and rates of oxygen consumption, and the authors suggested that this reduction in activity is a strategy to save energetic costs while deaminating free amino acids to stay iso-osmotic with the environment. To further investigate the time course of acute salinity transfer effects on oxygen consumption within our study, RMR was recalculated over hour-long intervals using the same data. In all groups acutely transferred from a higher to lower salinity there was a sharp initial increase in RMR in the first few hours followed by decreases towards baseline rates (Fig. 6). These patterns are consistent with a substantial cost of free amino acid catabolism, similar to the findings of Gilles (1973) and Yao et al. (2020). Interestingly, there were no such acute rises in metabolic rate upon temperature change (Fig. S3), bringing into question the relative costs of temperature and salinity changes in this species. It should be emphasized, however, that changes in activity are also occurring simultaneously with biochemical acclimatory changes, and their combined effects on metabolism cannot be disentangled without further research into the costs of free amino acid synthesis and catabolism in *T. californicus* in response to salinity change.

### Interactions between salinity and temperature and their consequences for the species

A main objective of this study was not only to describe the effects of salinity and heat independently on RMR and activity in *T. californicus*, but also to investigate the impacts of their interaction on sublethal measures of performance. Our findings show that salinity and heat have interactive effects, especially on copepod activity, and point towards possible ecological impacts of these stressors. In this splashpool species, decreased activity at high salinities and temperatures may inhibit effective foraging, predator avoidance, and mate-finding, potentially affecting the fitness of populations. Alternatively, high metabolic costs of low salinities in this genus may limit energy available for other activities, like somatic growth, reproductive investment, and biochemical mechanisms that promote survival under other stressors like high temperatures. This may be contributing to the previously documented decrease in acute heat tolerance in *T. californicus* at lower salinities (Denny and Dowd, 2022). In an experiment testing the effects of salinity and temperature on development in *T. brevicornis*, only copepods in normal seawater showed complete development, whereas copepods at low and high salinities had no survivorship past early nauplii stages or failed to produce clutches entirely (McAllen and Brennan, 2009). In that study, temperature was also found to affect reproductive characteristics like ovary development and clutch size. Additionally, *T. japonicus* cultured in salinities ranging from 4 to 32 ppt demonstrated greatest survival, growth rates, and fecundity in 32 ppt conditions relative to other treatments (Hagiwara et al., 1995). Further study of sublethal responses to salinity and temperature both individually and simultaneously may provide insight into the physiological and biochemical mechanisms underlying their interaction and higher-level fitness effects. Ultimately, the present work shows that changing temperature and salinity affect metabolic rates and rates of activity in the copepod *T. californicus*, and that these responses are difficult to disentangle. Research attempting to predict organismal or ecological responses using lab-based metabolic data, both in *Tigriopus* species and in other mobile aquatic organisms, should also consider activity rates within those models, as activity may have a strong influence on energy expenditure.

## Acknowledgements

Mike Phelps and Asaph Cousins provided feedback on experimental design. Mark Denny provided constructive comments on the manuscript.

## Data Availability

Upon publication, raw data will be available via the WSU Research Exchange (https://rex.libraries.wsu.edu/esploro/).

**Figure S1.**
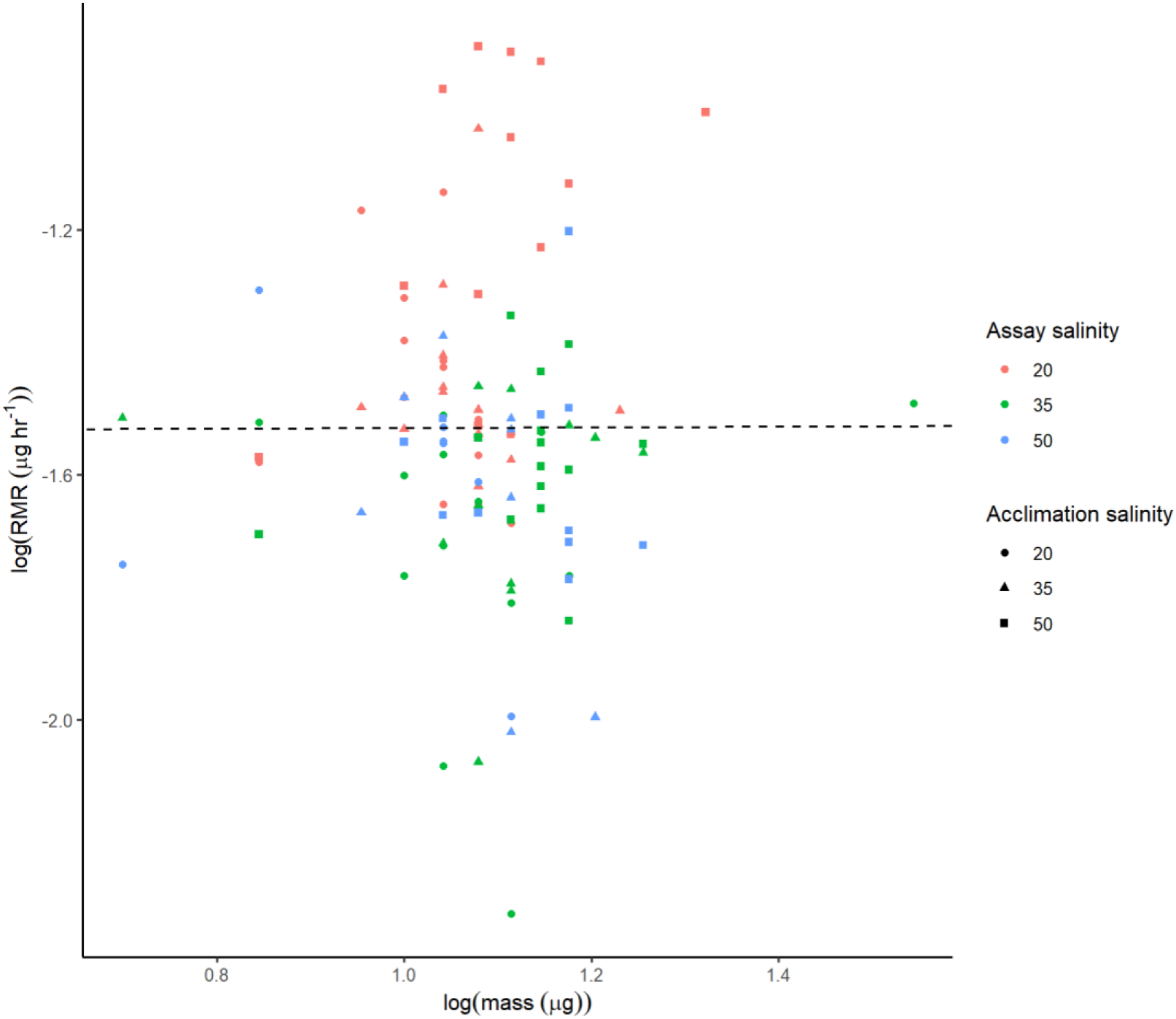
Log of copepod metabolic rates under 17.5°C does not scale allometrically with the log of mass. Dashed line represents the slope and intercept from a linear model (log(RMR) ∼ log(mass)) examining the effect of mass on metabolic rate of copepods measured under 17.5°C. Point color represents assay salinity, and point shape represents acclimation salinity. Because of the lack of allometric scaling and the unreliability of measured masses of 27.5°C copepods, we chose to forego correcting copepod RMRs by mass in our analyses.

**Figure S2.**
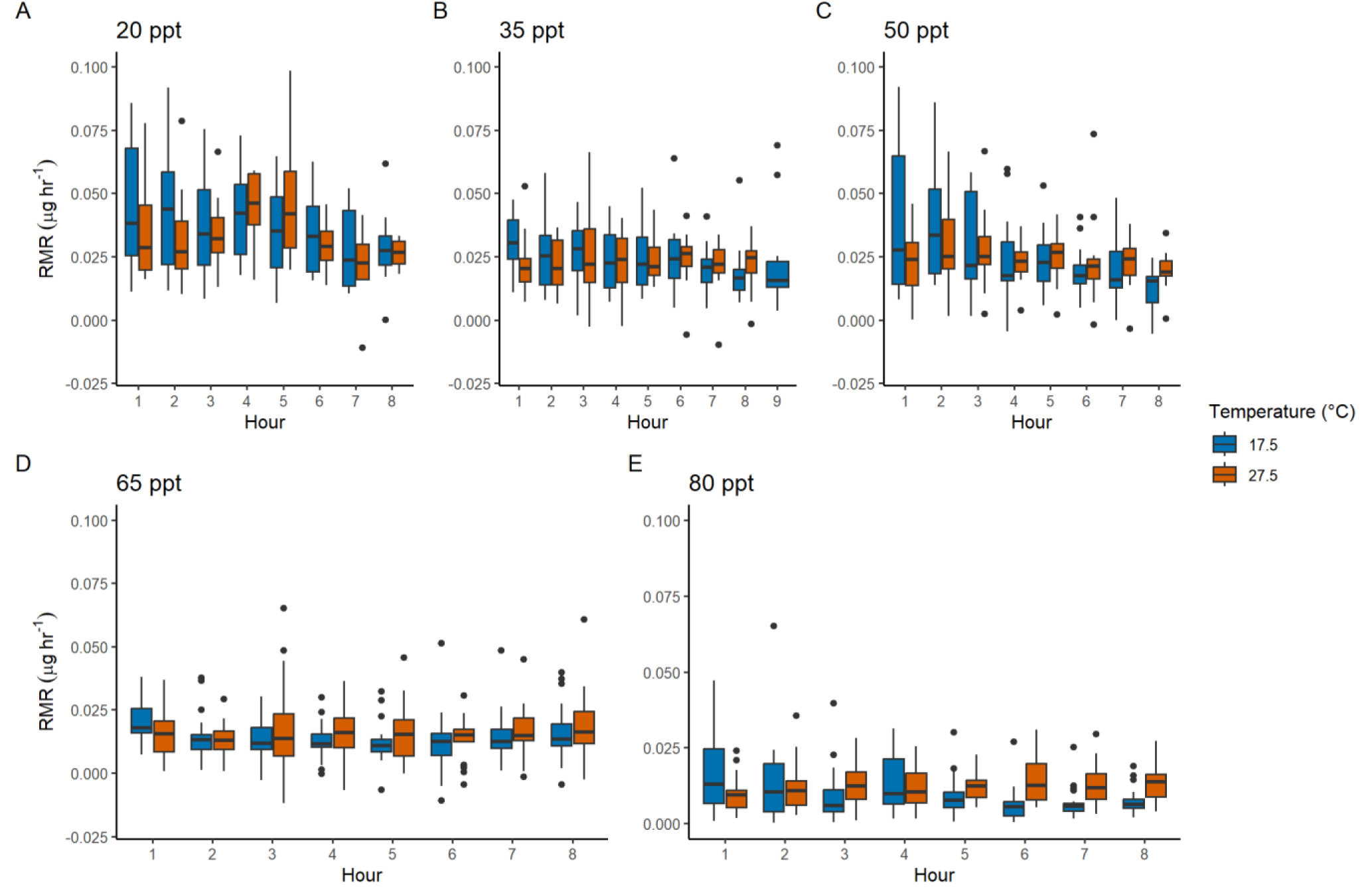
Hourly measures of RMR in chronic acclimation copepods show no acute temperature response. Based on the Arrhenius relationship we would anticipate 2-3x higher rates at 27.5°C than at 17.5°C, at least initially. Box plots representing RMR of copepods chronically acclimated to 20, 35, 50, 65, and 80 ppt (A-E, respectively) measured over hour-long periods. Median values are depicted by black lines in the middle of boxes, upper and lower box hinges correspond to the first and third quartiles, and whiskers extend to no further than 1.5 times the interquartile range.

**Figure S3.**
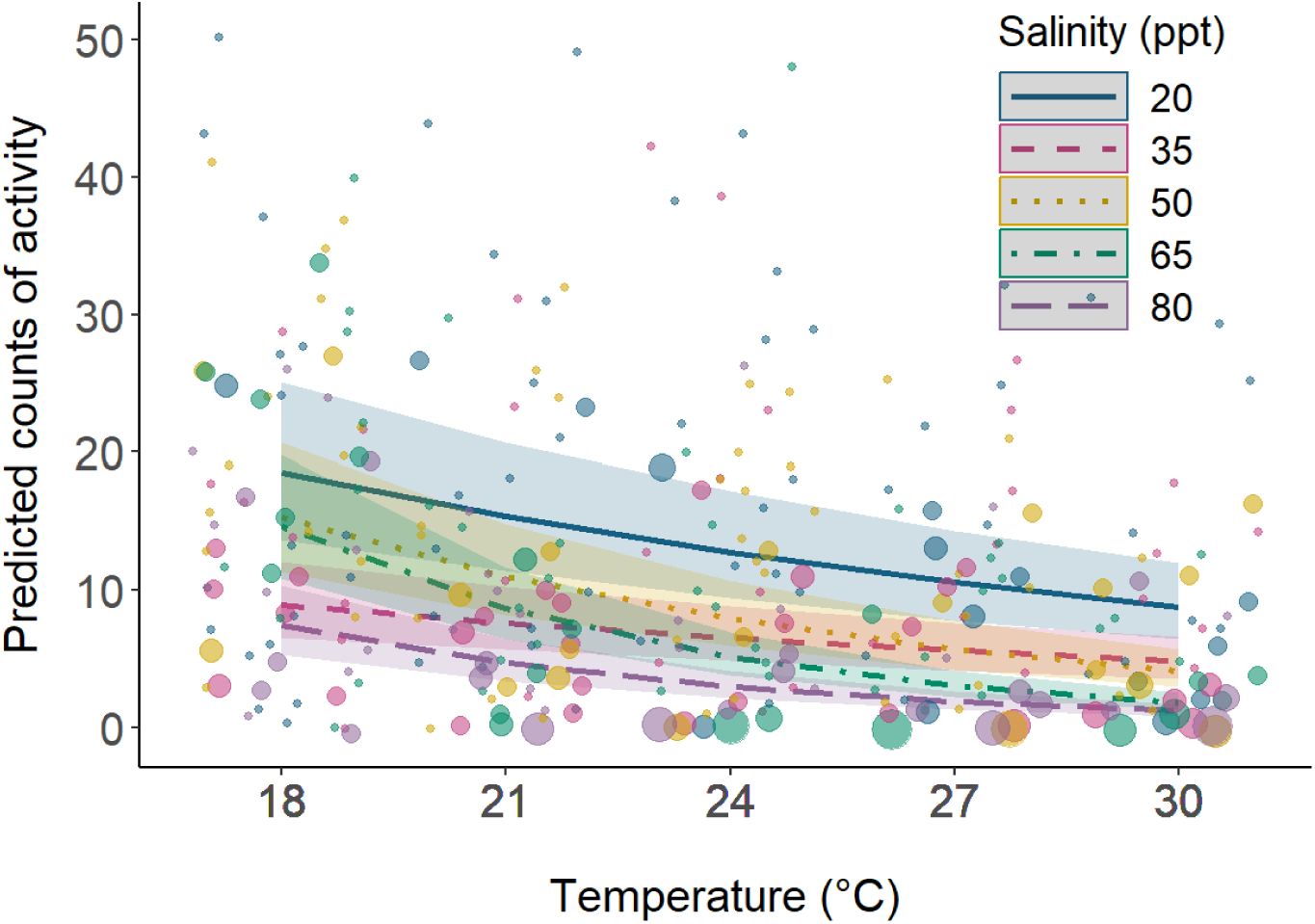
Copepod activity declines with increasing salinity and temperature. Predicted counts of activity are from a generalized linear mixed model with fixed effects of salinity, temperature, and an interaction between the two, and a random effect of individual. Colored ribbon around each line denotes the confidence interval for predictions at that salinity and temperature. Dots represent raw counts of activity, with size increasing with number of occurrences at that point.

**Figure S4.**
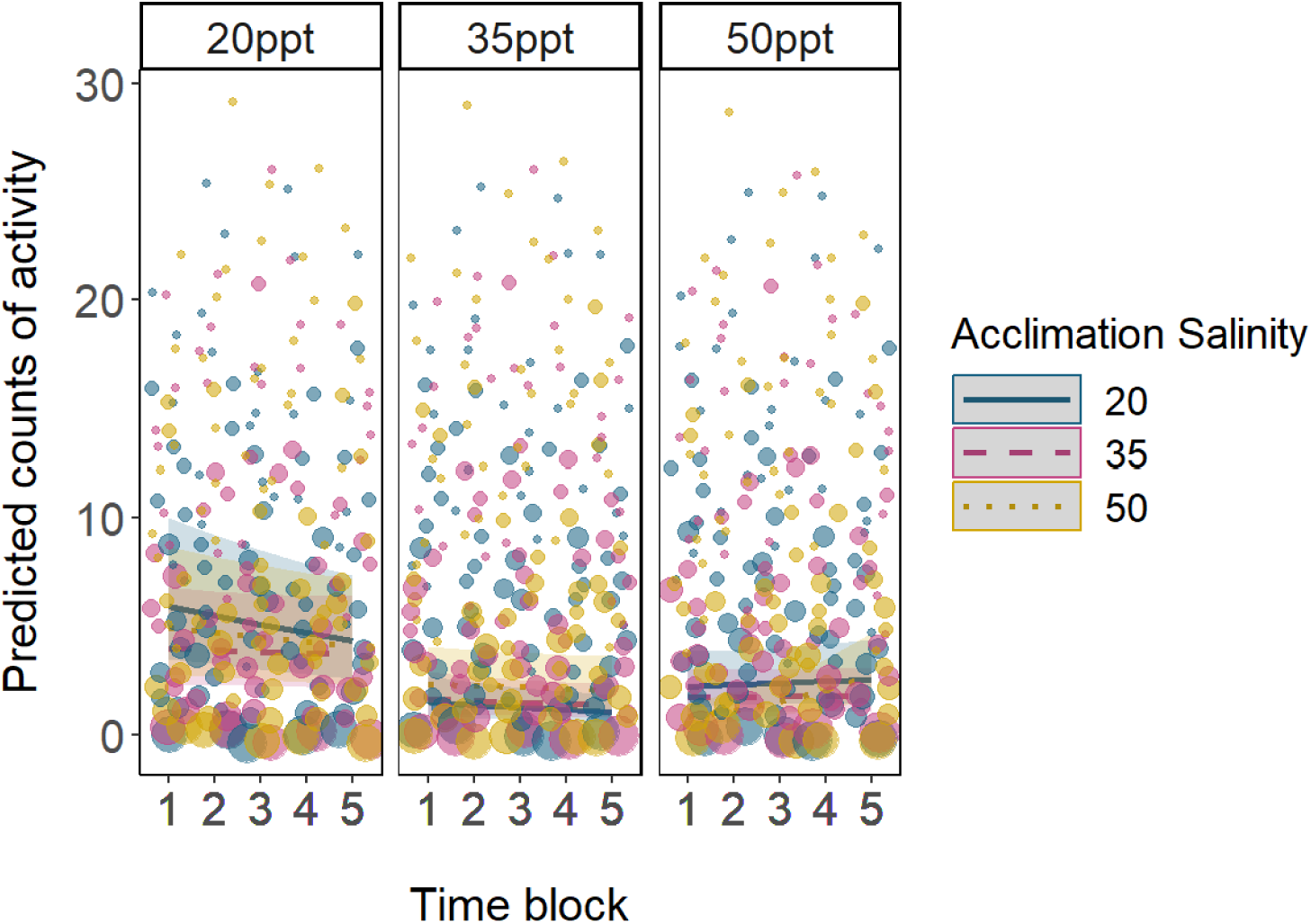
Acute salinity transfer impacts rates of copepod activity. Predicted counts of activity are from a generalized linear mixed model with fixed effects of acclimation salinity, assay salinity, time after transfer, interactions between all fixed effects, and a random effect of individual. Colored ribbon around each line denotes the confidence interval for predictions at that salinity and temperature. Assay salinities are represented by plot facets. Dots represent raw counts of activity, with size increasing with number of occurrences at that point.

**Table S1.** Summary of the statistical linear mixed effects model (with individual as a random factor) results testing for effects of chronic salinity acclimation, temperature, measurement interval, and their interaction on hourly copepod RMR.

| Factor | Chi square value | Df | p-value |
| --- | --- | --- | --- |
| **Salinity** | **72.8715** | **4** | **5.616e-15** |
| Temperature | 3.8032 | 1 | 0.051156 |
| **Hour** | **18.3611** | **8** | **0.018677** |
| Salinity*Temperature | 3.8546 | 4 | 0.426045 |
| **Salinity\*Hour** | **44.1159** | **28** | **0.027060** |
| **Temperature\*Hour** | **23.9016** | **7** | **0.001186** |
| Salinity*Temperature*Hour | 24.5297 | 28 | 0.653295 |

Significant terms are bolded. Measurements in this analysis were taken over the course of 9 hours and intervals (hour) consist of hour-long blocks. The model also included random effects of plate and copepod ID.

**Table S2.** Summary of the statistical linear mixed effects model (with individual as a random factor) results testing for effects of acute salinity transfer, temperature, measurement interval, and their interaction on hourly copepod RMR.

| Factor | Chi square value | Df | p-value |
| --- | --- | --- | --- |
| **Acclimation salinity** | **71.3384** | **2** | **3.229e-16** |
| Assay salinity | 5.7741 | 2 | 0.0557411 |
| Temperature | 2.3166 | 1 | 0.1279965 |
| Hour | 12.3208 | 8 | 0.1374554 |
| **Acclimation salinity\*Assay salinity** | **34.2080** | **2** | **6.755e-07** |
| **Acclimation salinity\*Temperature** | **14.2569** | **2** | **0.0008020** |
| Assay salinity*Temperature | 1.9763 | 2 | 0.3722603 |
| **Acclimation salinity\*Hour** | **90.9532** | **16** | **1.669e-12** |
| Assay salinity*Hour | 6.4151 | 14 | 0.9549207 |
| **Temperature\*Hour** | **16.3106** | **7** | **0.0224252** |
| Acclimation salinity*Assay salinity*Temperature | 6.2013 | 4 | 0.1846118 |
| **Acclimation salinity\*Assay salinity\*Hour** | **60.4167** | **28** | **0.0003602** |
| Acclimation salinity*Temperature*Hour | 20.8482 | 14 | 0.1055864 |
| Assay salinity*Temperature*Hour | 11.2698 | 14 | 0.6647226 |
| Acclimation salinity*Assay salinity*Temperature*Hour | 19.9473 | 28 | 0.8663773 |

Significant terms are bolded. Measurements in this analysis were taken over the course of 9 hours and intervals (hour) consist of hour-long blocks. The model also included random effects of plate and copepod ID.

